# The neonicotinoid acetamiprid is highly toxic to wild non-target insects

**DOI:** 10.1101/2024.04.11.589033

**Authors:** Jan Erik Sedlmeier, Ingo Grass, Prasanth Bendalam, Birgit Höglinger, Frank Walker, Daniel Gerhard, Hans-Peter Piepho, Carsten A. Brühl, Georg Petschenka

## Abstract

Although pesticides are seen as one of the main drivers of insect decline, there are still only few studies that assess their effects on non-target species under field conditions. Here we investigated the effects of the neonicotinoid insecticide Mospilan®SG (active ingredient acetamiprid) on plant bugs (Heteroptera: Miridae), a dominant group of European grassland insect communities. Abundance of three focal mirid species was reduced by up to 78% two days after field exposure at concentrations expected at field margins, with mortality varying considerably among species. Follow-up feeding assays with insecticide-treated host plants in the greenhouse and controlled dose-response assays in the laboratory confirmed the strong negative effects on non-target species. Strikingly, the neonicotinoid was nearly 10,000 times more toxic to plant bugs than to honeybees. In addition, male bugs were 20 times more sensitive than females in two of the three tested species. Thus, continuous exposure to neonicotinoids in the field may reduce the plant bug populations and promote a shift towards more insecticide-tolerant species, altering community composition. We suggest that sex-specific sensitivity should be considered in risk assessment and conclude that the real risk to non-target insects is currently greatly underestimated.

## 1. Main

We are facing an unprecedented decline in biodiversity, which is particularly evident for insects in the past years, as many studies have shown worldwide declines in insect biomass, abundance, and richness (Dirzo et al. 2014, Hallmann et al. 2017, IPBES 2019, Cardoso et al. 2020, Wagner et al. 2021). The intensive use of agrochemicals is considered to be a major driver of insect decline (Brühl and Zaller 2019, Wagner et al. 2021). Non-target insects can be exposed to insecticides in both agricultural and non-agricultural areas. For terrestrial insects, direct contact and oral uptake are likely the main routes of exposure. Direct contact can occur through overspray or contact with plant or soil surfaces containing residues. Oral uptake can occur through consumption of contaminated food, e.g. plants or other invertebrates (EFSA 2015). In terrestrial non-target habitats (e.g. field margins, neighbouring meadows, hedges), such exposure is mainly caused by overspray and spray drift (Schmitz et al. 2014). Spray drift is caused by wind during insecticide application and can contaminate habitats up to 100 m away from the field (van de Zande et al. 2015). Consequently, areas close to the field such as field margins, are particularly affected, since the first metre of a field margin is exposed to both overspray and spray drift. Estimates of pesticide deposition here range from 30% to 58% of the field rate (Schmitz et al. 2014, van de Zande et al. 2015). At the same time, semi-natural habitats such as field margins make up a significant proportion of insect habitats in modern agro-ecosystems (Hahn et al. 2014). It is therefore all the more alarming that there is increasing evidence that modern agro-ecosystems, as well as non-agricultural landscapes and even insects in nature preserves, are highly contaminated with pesticides (Brühl et al. 2021, 2024, Sabzevari and Hofman 2022).

While pesticide exposure of non-target insects seems quite obvious, the effect is still poorly understood. Although modern insecticides are sometimes claimed to be less toxic to important beneficial insects such as the Western honeybee *Apis mellifera* than to the target pest (e.g. based on the German bee risk levels B1-B4), sensitivity to insecticides can vary even between closely related species and is generally difficult to predict (Sánchez-Bayo 2012, Cabrera et al. 2017). The current EU insecticide registration protocol requires sensitivity testing for a limited number of non-target insects. These are typically the honeybee, parasitoid wasps, predatory mites, and individual representatives of large beetle families such as the Coccinellidae, Carabidae or Staphylinidae (Main et al. 2018). Furthermore, the vast majority of available ecotoxicological studies involve either aquatic invertebrates or terrestrial beneficial insects such as pollinators or predators (Teder and Knapp 2019, Rumschlag et al. 2020, Siviter and Muth 2020). Herbivorous insects, which account for about 50% of the total insect richness, have been largely neglected (Pisa et al. 2015). Another shortcoming in the ecotoxicology literature is the underrepresentation of community-level studies, i.e. evaluating a range of species with different sensitivities and effects on changes in predator prey relationships. The study of community-level effects in realistic exposure scenarios is an important cornerstone for assessing the true ecological consequences of insecticide contamination. Consistent community structure effects of insecticide contamination have already been shown for aquatic systems (Rumschlag et al. 2020). However, similar studies are lacking for terrestrial systems.

In this study, we address this research gap on insecticide exposure of non-target herbivorous insects, focusing on three main aspects: (1) realistic exposure scenarios, (2) community-level effects, and (3) differential sensitivity between closely related species and between sexes of the same species. We chose plant bugs (Heteroptera: Miridae) as a model group because they are a highly diverse family within the Heteroptera and a common component of non-target insect communities in agroecosystems (Wachmann et al. 2004). Plant bugs are highly abundant in meadows, but can also be found in grassy field margins, as many species are associated with Poaceae (Denys and Tscharntke 2002, Wachmann et al. 2004). These species feed on the panicles at the top of grasses, making them even more vulnerable to insecticide deposition than ground-dwelling insects (Kühne et al. 2002). However, this group of insects has been overlooked in previous studies on non-target effects of insecticides and is not considered in the risk assessment procedure for plant protection products. We argue that plant bugs are not only an ecologically important group to study, as true bugs make up a significant proportion of many bird diets (Exernová et al. 2003), but also provide an opportunity to investigate community-level effects and potential differences in sensitivity between species.

We conducted field, greenhouse, and laboratory experiments to mimic insecticide exposure in different scenarios. We used the neurotoxic neonicotinoid insecticide Mospilan®SG (Cheminova Deutschland GmbH & Co. KG, Stade, Germany) with the active ingredient acetamiprid in all experiments. Mospilan®SG is applied by spraying and is used in field crops such as rapeseed and potato, in orchards, in wine growing and in floriculture, in particular against chewing sucking pests. The active substance acetamiprid is both a contact and a systemic insecticide, as it can also be absorbed by plants and distributed throughout their tissues. Among other neonicotinoids, acetamiprid is used worldwide, but in the European Union it is the only neonicotinoid still registered for open-field use. In 2022, sales of the active ingredient in Germany alone amounted to almost 28 tonnes (BVL 2023).

In the field experiment, we sprayed three concentrations of Mospilan®SG in semi-open field plots with wild plant bug populations to simulate in-field and off-field exposure scenarios (Fig. 1A). The field plots were located in an extensively managed meadow in southern Germany. We selected treatments based on deposition estimates as follows: ’100%’ = maximum field rate; ’30%’ = exposure estimate for 1 m field distance (overspray + spray drift); ’0.15%’ = estimate for 20 m field distance (spray drift) (Fig. 1B). Plant bugs were sampled at 2, 9, 16, 24 and 30 days after application using a sweep net. We evaluated the total number of individuals of mirid populations within the field plots and conducted species-level analyses focusing on the three most abundant species: *Stenotus binotatus* (Fabricius, 1794), *Leptopterna dolabrata* (Linnaeus, 1758), and *Megaloceroea recticornis* (Geoffroy, 1785). These three species overlap in phenology (May – August) and host plants (Poaceae), and also differ in size (on average 6.6 mm *S*. *b*., 8.3 mm *L*. *d*., and 9.1 mm *M*. *r*.): We also conducted pesticide analyses of vegetation samples from the plots to monitor insecticide exposure and metabolism in plants over time.

**Figure 1:**
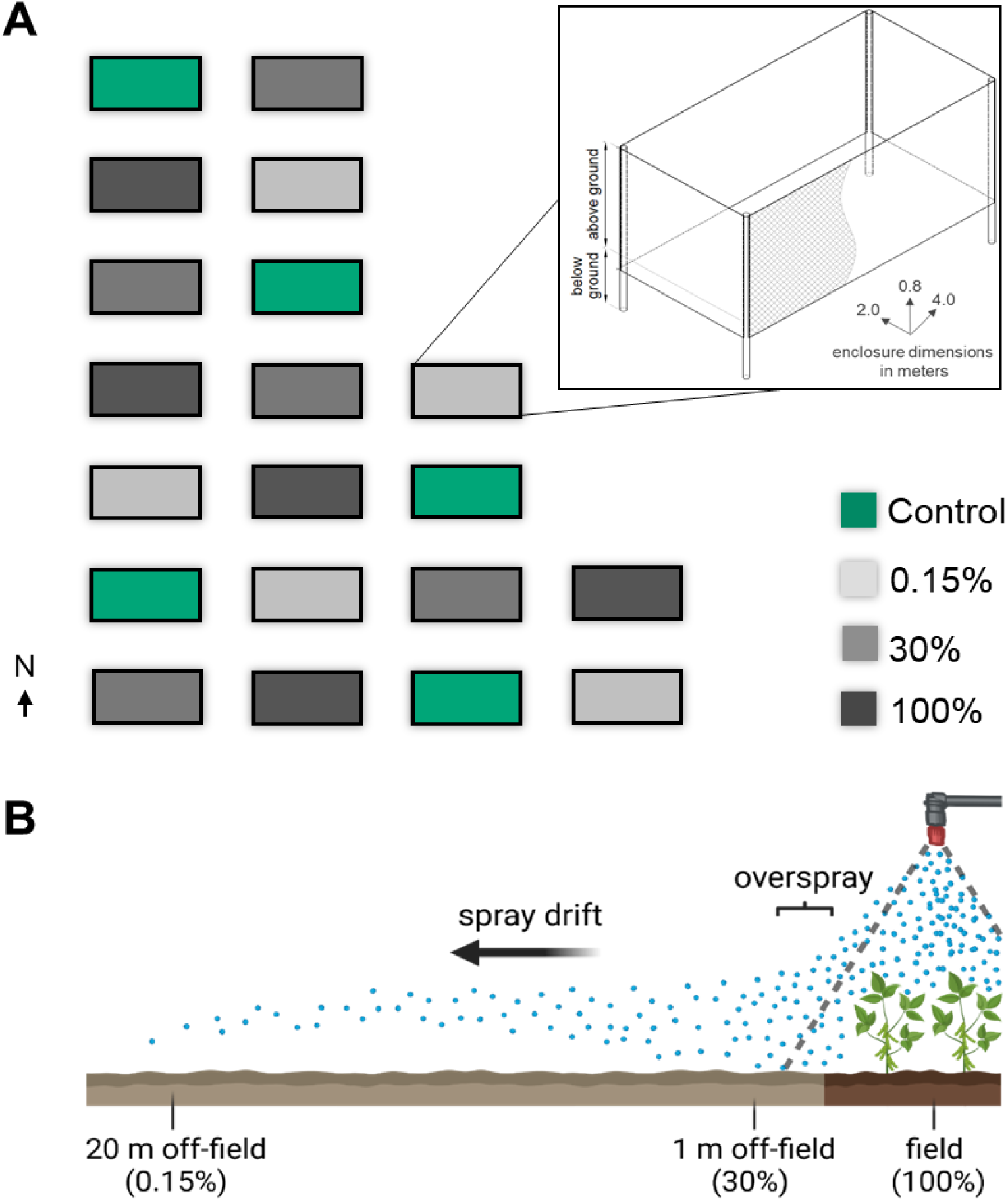
Overview of the field experiment and concept of treatment choice. (A) Field enclosures are separated by 2 m. Coloration indicates Mospilan®SG – treatment based on contamination estimates. Drawings created with AUTODESK AutoCAD 2023 and Fusion 360. (B) Treatment choice (schematic drawing) is based on literature estimates: 100% = maximum field rate, 30% = contamination estimate for 1 m field distance (overspray + spray drift, Schmitz et al. 2013), 0.15% = contamination estimate for 20 m field distance (spray drift, JKI 2020). Graph in panel B created with BioRender.com.

We further assessed insecticide susceptibility in a greenhouse experiment using host plants (Poaceae from the same meadow) that were sprayed with Mospilan®SG two days earlier to simulate an indirect exposure, where non-target insects enter a habitat after insecticide deposition. In this experiment, we recorded the survival of the three focal species after four days of exposure (i.e., after 6 days of plant exposure). We also analysed the insecticide exposure of the host plants two days after application. Finally, we performed dose-response assays in the laboratory, where we applied different concentrations of Mospilan®SG dorsally to individual plant bugs (see Methods) and recorded mortality after 24 hours. This approach allowed us to investigate intrinsic differences in contact sensitivity between species and between conspecific males and females without confounding factors, such as potential behavioural avoidance of exposure.

## 2. Results

### 2.1. Field experiment

Neonicotinoid application reduced mirid abundance by 83% (100% treatment) and 82% (30% treatment) after two days compared to the control (100%: p<0.0001; 30%: p<0.0001; Tukey-test) (Fig. 2A). Nine days after exposure, the abundance was reduced by 70% (100% treatment) and 57% (30% treatment) compared to the control (100%: p<0.0001; 30%: p<0.001; Tukey-test). This pattern was less pronounced in later samples, while the total plant bug population declined in all treatments, including the control, presumably due to plant bug phenology. No difference in mean counts was observed between the control and 0.15% treatments and between the 30% and 100% treatments at any sampling event.

**Figure 2:**
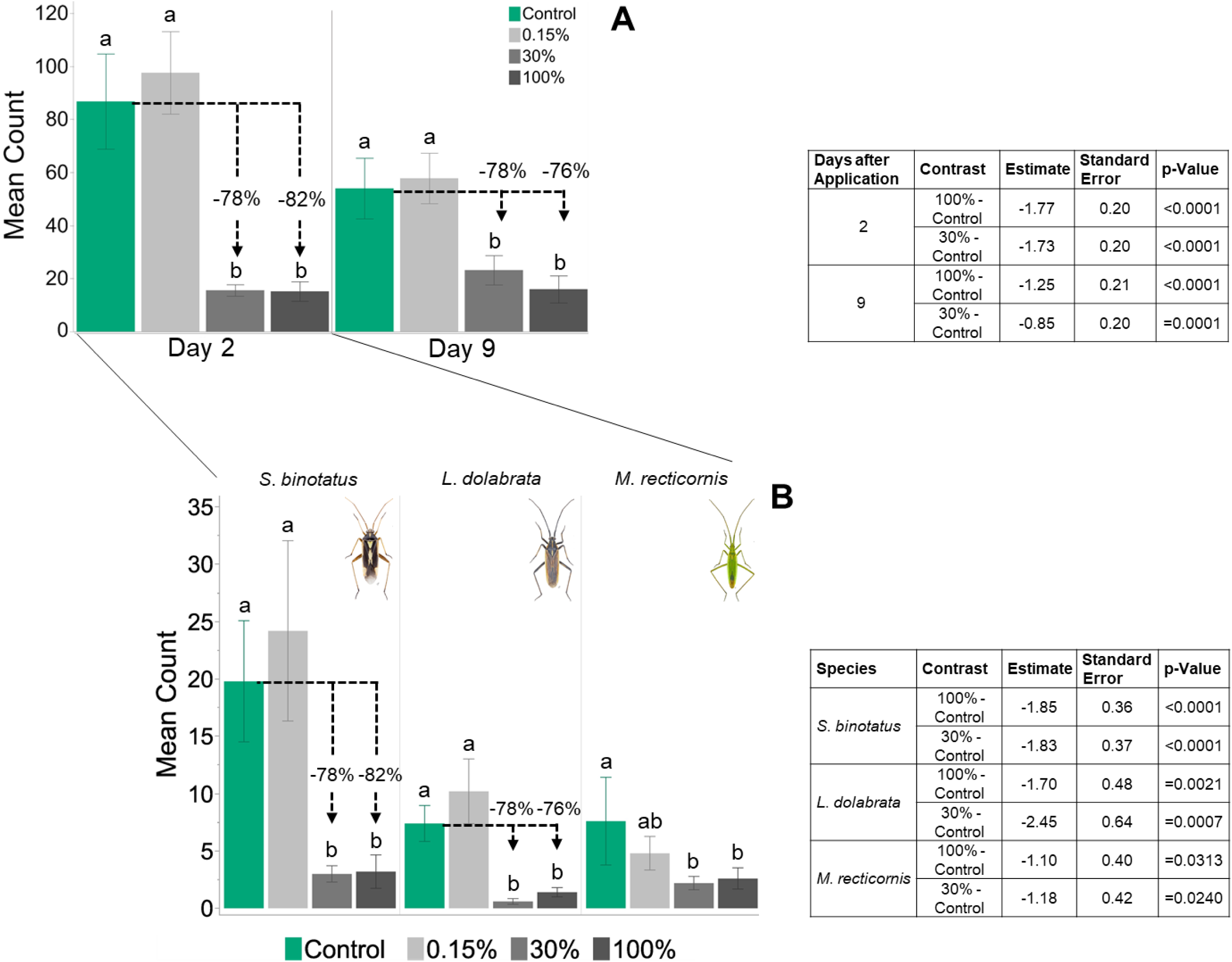
Exposure to Mospilan®SG reduces plant bug abundance and causes a species-specific response. A: Mean number of imagines and nymphs two- and nine days after insecticide application (± standard error). B: Mean number of imagines two days after insecticide application separated by species (± standard error). *Stenotus binotatus* (left), *Leptopterna dolabrata* (center) and *Megaloceroea recticornis* (right). Means within the same sampling day (A) or the same species (B) not sharing the same letter are significantly different (Tukey-test at the 5% level of significance). Results for significant contrasts are provided in the adjoining tables. Numbers above bars indicate percentage difference to control (rounded numbers); n = 5 for all treatments and species (panel A + B). Insect images © Gerhard Strauss, Biberach.

Focusing on the three most abundant species, the mean counts for *S*. *binotatus*, *L*. *dolabrata* and *M*. *recticornis* in the 30% and 100% treatment were significantly reduced compared to the control two days after insecticide application (*S*. *b*. 100%: p<0.0001, *S*. *b*. 30%: p<0.0001; *L*. *d*. 100%: p=0.0021; *L*. *d*. 30%: p<0.001; *M*. *r*. 100%: p=0.031; M. r. 30%: p=0.024) (Fig. 2B). This pattern partially shifted with later samplings, however, in line with the known phenologies the populations drastically declined in later samplings, including the control (Fig. 3).

**Figure 3:**
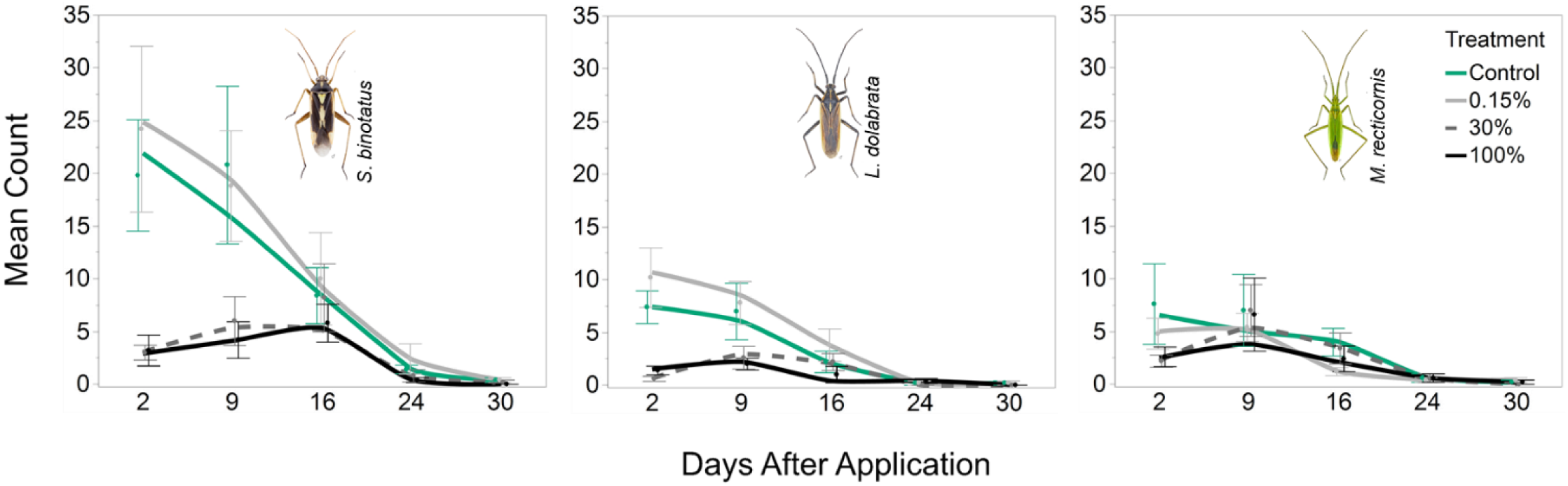
**Population development in the field trial** for *Stenotus binotatus* (left), *Leptopterna dolabrata* (center) and *Megaloceroea recticornis* (right). Curves are fitted with the Kernel method (local fit: linear). Mean number of imagines per treatment (± standard error); n = 5 for all treatments and samplings. Insect images © Gerhard Strauss, Biberach.

Pesticide analysis of vegetation samples from the enclosures taken after 7-, 17- and 30 days revealed residues of acetamiprid and its metabolite acetamiprid-N-desmethyl in the 100% and the 30% treatments (Table 1). No residues were found in the 0.15% treatment or in the control at any sampling time. Further information on pesticide analysis is provided in supplementary file 1.

**Table 1:**
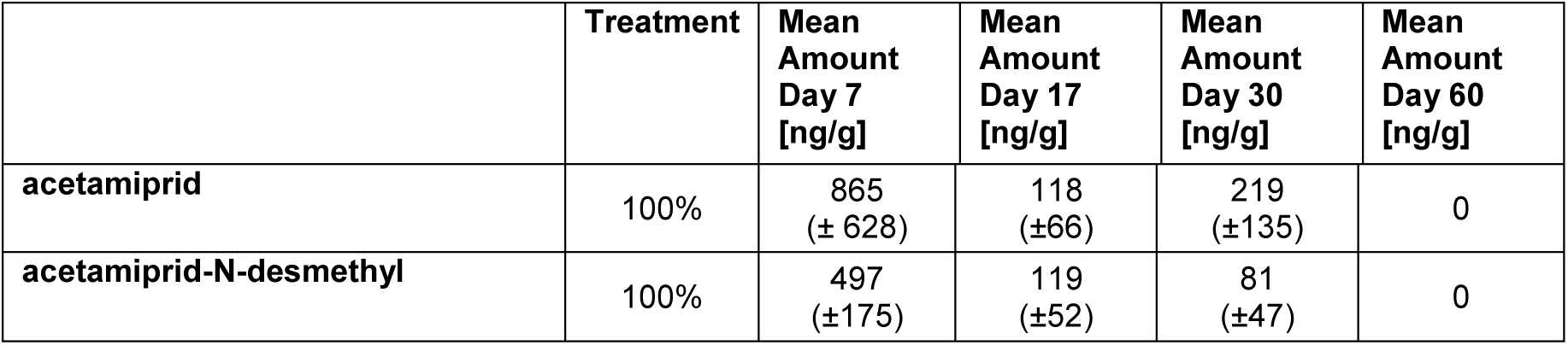
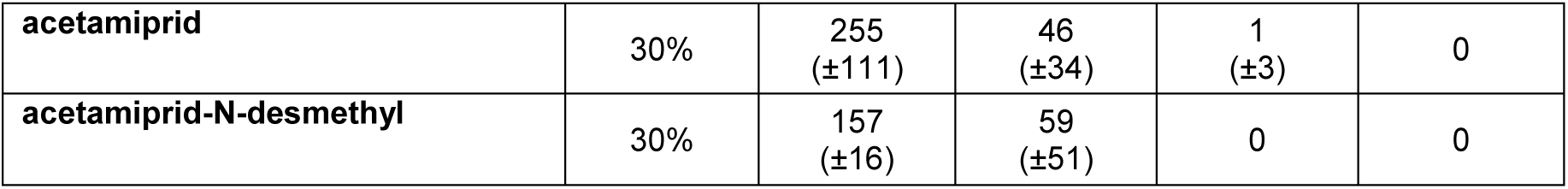
Mean concentrations of acetamiprid and the metabolite acetamiprid-N-desmethyl (± SD) per gram dry mass of plant tissue sampled inside the 100% and 30% field enclosures (LC-MS) after application of Mospilan®SG in the field experiment. Numbers are rounded, n = 5 for each treatment and substance.

|  | Treatment | Mean Amount Day 7 [ng/g] | Mean Amount Day 17 [ng/g] | Mean Amount Day 30 [ng/g] | Mean Amount Day 60 [ng/g] |
| --- | --- | --- | --- | --- | --- |
| acetamiprid | 100% | 865 (± 628) | 118 (± 66) | 219 (± 135) | 0 |
| acetamiprid-N-desmethyl | 100% | 497 (± 175) | 119 (± 52) | 81 (± 47) | 0 |
| acetamiprid | 30% | 255<br>(±111) | 46<br>(±34) | 1<br>(±3) | 0 |
| acetamiprid-N-desmethyl | 30% | 157<br>(±16) | 59<br>(±51) | 0 | 0 |

### 2.2. Greenhouse experiment

Survival of the three plant bug species four days after being placed on plants sprayed 48 h before with 30% of the maximum field application rate of Mospilan®SG was reduced in all species. Specifically, survival was reduced by 88% for *S*. *binotatus*, 39% for *L*. *dolabrata*, and 35% for *M*. *recticornis* compared to the respective controls (Fig. 4). In none of the nine replicates did any individuals of *S*. *binotatus* survive the insecticide treatment.

**Figure 4:**
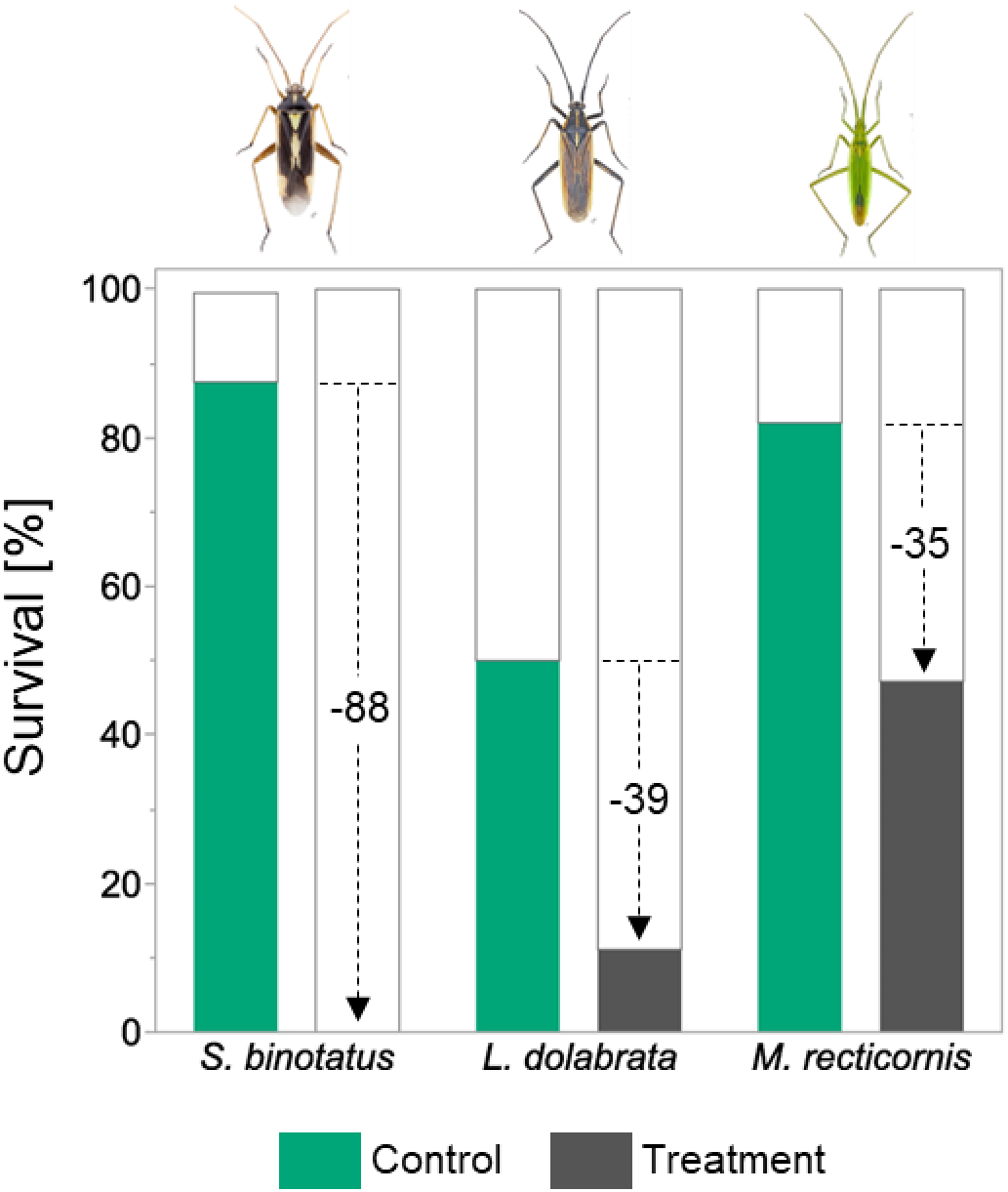
Survival of mirids after four days of exposure to plants treated with 30%- Mospilan®SG. Plants were sprayed with insecticide two days prior to plant bug release. Survival rates at day four are given for *Stenotus binotatus*, *Leptopterna dolabrata* and *Megaloceroea recticornis*. Numbers within bars indicate difference between control survival and treatment survival; n = 9. Insect images © Gerhard Strauss, Biberach.

**Figure 4:**
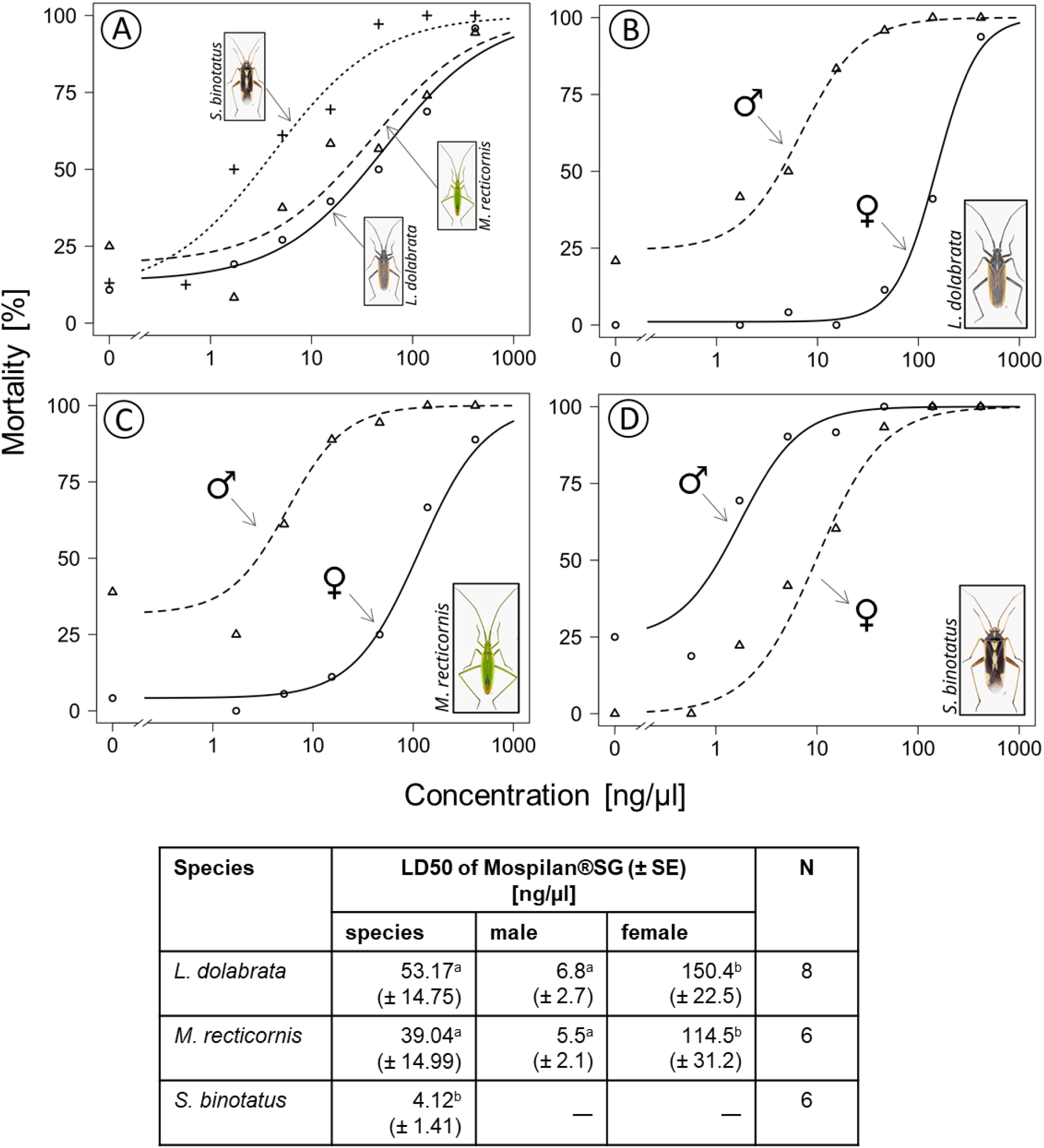
Mospilan®SG dose-response curves for all species (A) and separated by sex for *Leptopterna dolabrata* (B), *Megaloceroea recticornis* (C) and *Stenotus binotatus* (D). Data points represent mean values per concentration. Derived LD_50_ values are presented in the table below. Separate LD_50_ values for male and female specimens were generated only for those species in which the sensitivity to the insecticide was significantly different between the sexes based on the fitted generalised linear mixed model (GLMM). LD_50_ values of species not sharing a common letter are significantly different. LD_50_ values were compared using the compParm function of the ‘drc’ package in R. LD_50_ values of males and females of the same species not sharing a common letter are significantly different. Abbreviations: SE = standard error; N = number of biological replicates (i.e., petri dishes) per concentration. Total number of specimens tested: 245 for *S*. *binotatus*, 332 for *L*. *dolabrata*, and 224 for *M*. *recticornis*. Note that mean values for the lowest Mospilan®SG treatment for males and females of *M*. *r*. (C) consist of only two data points instead of three. Insect images © Gerhard Strauss, Biberach.

Acetamiprid and the metabolite acetamiprid-N-desmethyl were found in all samples collected from treated plants two days after application (i.e. the time of plant bug release). The mean concentration was 267 (±78) ng/g plant material for acetamiprid and 62 (±33) ng/g for acetamiprid-N-desmethyl (dry weight; n = 9; rounded numbers). Further information on pesticide analysis is provided in supplementary file 1.

### 2.3. Dose-Response assays

All three species showed high sensitivity to the Mospilan®SG treatment in the dose-response assays in the laboratory (Fig. 4). The observed LD_50_ values correspond to only 12.8% of the maximum field rate for *L*. *dolabrata*, 9.4% for *M*. *recticornis*, and 1.0% for *S*. *binotatus*. In addition, species-specific differences were observed, with *S*. *binotatus* being significantly more sensitive to the insecticide than the other two species based on LD_50_ comparisons (*L*. *dolabrata* – *S*. *binotatus*: p<0.01, t=3.33; *M*. *recticornis* – *S*. *binotatus*: p=0.02, t=2.33).

Besides a very high overall toxicity of Mospilan®SG, we observed significant differences between males and females of two species. Based on LD_50_ values, males of *L*. *dolabrata* and *M*. *recticornis* were >20 times more susceptible than females (*L. d.* t=6.33, p<0.0001; *M. r.* t=3.48, p=0.0005). There was no difference between sexes for *S*. *binotatus* based on the fitted GLMM prior to the dose-response analysis.

## 3. Discussion

Insecticides are a prime suspect in insect declines, but there is a lack of empirical evidence of real-world effects on non-target species outside the cropping area. To understand the impact of terrestrial insecticide contamination on non-target insect communities, we conducted a series of field, greenhouse, and laboratory experiments to investigate community-, species- and sex-specific responses in the model insect group of mirid bugs. Mirid bug species in the field experiment were highly sensitive to neonicotinoid exposure. We found species-specific responses across different exposure scenarios and drastic differences in sensitivity between males and female of two species tested in the laboratory.

### 3.1. Mirid communities are highly sensitive to insecticide contamination

Consistent across experiments, our study shows a high susceptibility of mirids to insecticide exposure. In the field experiment, we aimed to simulate a scenario of direct drift contamination with specimens present in the respective off-field environment during spray application. A drastic reduction in mirid abundance was observed in the scenario representing exposure within one metre of the field (30% of maximum field rate).

A second scenario of indirect drift contamination for the same field distance was simulated in the greenhouse experiment, as it is also possible that insects enter an area only after drift contamination occurred (i.e., after insecticides were applied). Residues of the compound that may still be present on surfaces (soil or plant material) hours or days later can still be absorbed through an insect’s cuticle, e.g. via the tarsi (Yu 2015), or ingested during feeding. A special case, however, is that of systemic insecticides, where the active ingredient can also be taken up by host plants of non-target insects, providing a route for oral contamination of piercing sucking herbivorous insects (e.g., mirids) for presumably even longer periods of time. In our experiment, even two days after host plants were treated with 30% of the maximum field rate of Mospilan®SG, the survival rates of mirids placed on these plants were significantly reduced.

These results clearly demonstrate a detrimental effect of insecticide contamination on non-target mirid populations located close to arable fields. Further, a generally high sensitivity to neonicotinoid insecticides of this ecologically important insect group that occupies an important position in the trophic pyramid, is revealed (Exernová et al. 2003). This was further supported by the insecticide dose-response assays. Here, LD_50_ values for three mirid species were at concentrations as low as 1% of the original field rate. To put this difference into context, these values can be compared to those of common model organisms such as the Western honeybee *A. mellifera*. However, LD_50_ values from dose-response assays are typically given as a concentration of active ingredient per microliter or per individual. To establish comparability, the values we obtained using a commercial insecticide formulation can be adjusted to reflect the 20% active ingredient acetamiprid content. This corresponds to 10.63 ng/bug for *L*. *dolabrata*, 7.81 ng/bug for *M*. *recticornis* and 0.82 ng/bug for *S*. *binotatus*. For honeybees an LD_50_ of ∼ 8090 ng active ingredient per bee is reported as contact exposure value (European Commission 2004). Accordingly, the sensitivity of the tested mirid species is ∼ 760 times (*L*. *dolabrata*) to ∼ 9866 times (*S*. *binotatus*) higher.

However, the toxicity of a pure active ingredient may differ from that of a product formulation (Nagy et al. 2020). Therefore, such comparisons should be made with caution. Nevertheless, this huge difference in LD_50_ values shows that mirids, and probably other non-target insects, are very sensitive to neonicotinoid insecticides and consequently to environmental contamination at only a fraction of the applied field rates. This group of insects must therefore be given greater consideration in risk assessment (see 3.4 below).

### 3.2. Species-specific sensitivity

The three mirid species that were tested in this study can be considered as model species representing non-target phytophagous insects. In this context, these species are of particular interest because they share a similar ecology (e.g., feeding style, phenology and host plants, Wachmann et al. 2004) and yet showed markedly different insecticide susceptibility under the experimental contamination scenarios. Differences in species response were already suggested by the field experiment and further confirmed by the greenhouse experiment with treated plants and the dose-response assays in the laboratory. As differences in species response were not only observed in the field experiment, it is unlikely that they are related to feeding preferences or the spatial position of the bugs on the plants during spraying, but rather follow from species-specific physiological traits.

Physiological factors that have been proposed to influence sensitivity include, for example, differences in the activity and abundance of detoxification enzymes, or simply body size (Müller 2018). Notably, the smallest (and lightest) species *S*. *binotatus* was always the most sensitive in our greenhouse- and dose-response assays, suggesting that body size or mass is an important factor. Goulson (2013) compared available LD_50_ values from neonicotinoid studies (topical and oral) and stated that much of the variation in sensitivity between several insect taxa could be explained by differences in body size/weight. For example, the LD_50_ for the same neonicotinoid of the highly sensitive *Nilaparvata lugens* (body weight ∼1 mg) is similar to the one of the least sensitive *Leptinotarsa decemlineata* (body weight ∼ 130 mg) expressed as ng active ingredient per mg body weight (0.82 and 0.67). Considering the species we tested, although *L*. *dolabrata* has more than twice the body mass of *M*. *recticornis*, the LD_50_ values are almost identical when expressed as concentration of active ingredient per body mass (L. d.: 0.53, M. r.: 0.66; see supplementary file 2). At the same time, there is a more than 4-fould difference to the LD_50_ of *S*. *binotatus* (0.13). Therefore, body weight alone cannot explain the difference in sensitivity between these species.

While the physiological mechanisms involved remain to be investigated, differences in species susceptibility could permanently alter local community structure under a constant or repeated contamination scenario. Changes in invertebrate community composition have been associated with pesticide contamination in the past. However, this has mainly been studied in freshwater communities (Beketov et al. 2013, Rumschlag et al. 2020), whereas the effects on terrestrial communities are less well understood. A recent meta-analysis showed that pesticide contamination reduces soil invertebrate diversity (Beaumelle et al. 2023). Main et al. (2020) have also linked neonicotinoid soil contamination to reduced species richness of wild bees in field margins. More research is needed to understand the long-term consequences of pesticide exposure on terrestrial non-target invertebrate communities on a landscape scale. Particular attention should be paid to univoltine species with short phenologies that may overlap with common pesticide application regimes. Populations of such species, such as the mirids in this study, should be particularly limited to cope with pesticide contamination should it occur during their short and only reproductive phase throughout the year.

### 3.3. Sex-specific sensitivity

In addition to differences in sensitivity at the species level, we found that males of *L*. *dolabrata* and *M*. *recticornis* were significantly more sensitive to the insecticide than the corresponding females. Sex-specific insecticide sensitivity is a pattern reported for many insects, with males often being more susceptible than females (Huntzinger et al. 2008, Andreazza et al. 2020). However, there are also contrary reports in other species (Navarro-Roldán et al. 2017).

Potential factors involved in sex-specific susceptibility partly overlap with those likely driving the observed species-specific differences. The most likely factor is again body mass, especially in sexually dimorphic species such as *L*. *dolabrata* and *M*. *recticornis*. In both species, males had a lower mean fresh body mass than females, with a 2.9-fold difference in *L*. *dolabrata* and a 2.2-fold difference in *M*. *recticornis* (n = 10 per species and sex). This is not correlated with the difference in LD_50_ values between males and females, which is 22-fold for *L*. *dolabrata* and 21-fold for *M*. *recticornis*. However, such relationships may not be necessarily linear.

Another source for differential susceptibility is at the enzymatic level. Frohlich (1988) showed that the activity of detoxification enzymes decreased with age in males of the solitary bee *Megachile rotunda*, but not in females. An age-dependent change in enzyme activity was also observed by Guirgus and Brindley (1975) when they compared 1-day-old males of *Megachile pacifica* with 4-day-old males, both of which had been previously treated with a fungicide. Thus, age also appears to play a critical role in susceptibility of insects to insecticides.

The higher control mortality observed in males in our dose-response study (which was accounted for in the models) would suggest an older initial age compared to females in the same cohort. However, to our knowledge, there is no evidence of a sex-specific time lag in the emergence of adult specimens of these species, and according to our observations, males and females appeared simultaneously in the field. However, for *L*. *dolabrata*, Wachmann et al. (2004) suggest that females live longer than males. A lower vitality of male specimens due to a shorter life cycle could therefore explain a higher control mortality and further higher susceptibility to stressors such as an insecticide. Regardless of the underlying mechanisms, the observed differences could lead to a shift in sex ratios, in this case towards females. This could ultimately lead to a reduction in abundance and, in the long term, a shift in species composition.

### 3.4. Environmental risk assessment

To assess the risk of insecticide contamination to a non-target species, it is essential to determine the true vulnerability of that species in its environment. Vulnerability involves both physiological sensitivity and exposure to an insecticide (de Lange et al. 2012). We have shown that mirids, as a non-target group, are physiologically highly sensitive to insecticides. Furthermore, the species tested here are potentially found in highly exposed areas of the off-field environment, i.e. in the upper vegetation of field margins (Kühne et al. 2002, Wachmann et al. 2004). We also demonstrated that exposure to a systemic insecticide extends beyond the initial contamination, as plant bugs placed on previously sprayed plants showed a drastic reduction in survival. Considering that in the field experiment we were able to detect acetamiprid in host plant tissue up to 30 days after application, the threat to non-target plant bugs is clear. It is therefore all the more surprising that the current risk assessment does not consider this group or other herbivorous insects in the non-target insect community. In addition, field margins less than 3 m wide are not currently considered to be terrestrial non-target habitats for pesticide application (Kühne et al. 2000). Consequently, this habitat and the insects living in it are not protected from pesticide contamination (Hahn et al. 2014).

Roß-Nickoll et al. (2004) argue that the test procedures used in the EU risk assessment are insufficient because the test species used do not adequately reflect the non-target insect community. As a result, the disruption to these communities caused by pesticide contamination is not taken into account. This discrepancy is further illustrated by the drastic difference in LD_50_ values between the honeybee model and the mirid species we tested which suggests an up to 10,000x higher toxicity for mirids compared to the honeybee. There is also the problem that risk assessment is currently only done at the species level, and that sex-specific sensitivity is not considered. However, the up to 22-fold difference in LD50 values between sexes that we found could scale to detrimental long-term effects on community composition. This calls for an increase of the so-called uncertainty factors used to cover different species and sexes from currently 10 to more than 1000. Consequently, and in light of the renewal of the authorisation of acetamiprid until 2033 (European Commission 2018), we argue that the current European risk assessment system urgently needs to be adapted accordingly to adequately address the environmental risk of pesticide contamination.

## 4. Methods

### 4.1. Statistical Analysis and Graphical Representation

Statistical analysis was conducted using R (version 4.3.0, R Core Team 2023) and R-Studio (version 2023.6.1.524, Posit team 2023). Therefore, only specific packages are cited in the following sections.

All graphs showing bar or stacked bar charts were generated using JMP®Pro (version 17.2.0, SAS Institute Inc. 2023). Dose-response curves were generated with R using the package ‘drc’ (Ritz et al. 2015) and edited with Inkscape (version 1.3.2, Inkscape Project 2020).

### 4.2. Field Experiment

#### 4.2.1. Construction

In May 2022, we constructed 20 field plots (2 x 4 m) on an extensively managed meadow (48°42’31’’N 9°12’48’’E) in the botanical garden of the University of Hohenheim, Stuttgart, Germany. In every corner of each plot, a 1.4 m long wooden pole was vertically inserted 40 cm into the soil. An 85 cm high insect gauze (mesh size: 0.27 x 0.79 mm, *Howitec Netting bv*, AH Joure, the Netherlands) was vertically stretched around these poles. About 10 cm of the gauze was inserted into the soil to form a barrier, while 75 cm remained aboveground fixed to the poles. Thereby, the top of the field plot remained uncovered. These semi-open field plots (henceforth, enclosures) were additionally braced with tent cords that were attached at every corner. On an area of 340 m^2^, we set up 20 enclosures in seven rows with each being separated by the neighbor by 2 m on all sides. The field experiment was laid out as a randomized block design with four treatments: three different concentrations of insecticide and a control. Each enclosure was defined as an individual block with every treatment being replicated five times (Fig. 1). The choice of blocks was made with respect to the environmental conditions on the site. Due to surrounding trees, the duration of solar radiation was longest in the northwestern area and shortest in the southeastern area. In addition, the test site was located on a south-facing slope. This resulted in a soil moisture gradient towards the south slope. The following blocks were assigned accordingly: A = upper slope + high solar radiation, B = middle slope + high solar radiation, C = middle slope + medium solar radiation, D = lower slope + medium solar radiation, E = lower slope + low solar radiation (see supplementary file 3).

#### 4.2.2. Insecticide application

We used the nerve-acting neonicotinoid insecticide Mospilan®SG (FMC Agricultural Solutions, Cheminova Deutschland GmbH & Co. KG, PO Box 2047, Stade, Germany), with the active ingredient acetamiprid for all experiments described in this study.

The following concentrations were applied in the field experiment by spraying: (1) typical field application rate of 250 g/ha in 600 l of water (100%), (2) 30% of the field application rate, (3) 0.15% of the field application rate. These values are based on estimates for overspray- and drift contamination with respect to the field distance (Schmitz et al. 2014, JKI 2020). The underlying concept is illustrated in figure 1A. In the control, only water was applied. The application rate was scaled down to the field plot area of eight square meters (Tab. 2). The amount of 0.48 l containing the respective insecticide concentration was applied equally to each field plot using the *GLORIA prima 3* hand sprayer (GLORIA Haus-& Gartengeräte GmbH, Witten, Germany).

**Table 2:** Application rates of Mospilan®SG of the field experiment.

| Treatment | Application Rate | Application Rate in Field Plot | Stock Solution |
| --- | --- | --- | --- |
| 100% | 250 g/ha in 600 l water | 200 mg/8 m<sup>2</sup> in 0.48 l water | 1.25 g in 3 l water |
| 30% | 75 g/ha in 600 l water | 60 mg/8 m<sup>2</sup> in 0.48 l water | 0.375 g in 3 l water |
| 0.15% | 0.375 g/ha in 600 l water | 0.3 mg/8 m<sup>2</sup> in 0.48 l water | 3.125 mg in 5 l water |

The hand compression sprayer was calibrated before use and the volume output was measured. The mean deviation of liquid output was around 1%. Details of the calibration procedure are given in supplementary file 4.

Application started with the control (water only) followed by the insecticide treatments in increasing concentration. Between treatments, the hand sprayer was completely emptied and re-calibrated. To control for uneven application due to a change in volumetric flow rate (because of decreasing pressure during spraying), the field plots were divided into four equal quarters. Each quarter was assigned a number from one to four. Spraying would always start in the quarter numbered one and end in the quarter numbered four. The order of numbers was randomized within plots of the same treatment.

During application inside the plots, the spray gun was opened with the nozzle pointing downwards towards the soil while moving in a standardized pattern of vertical and horizontal trajectories at a height of 80 cm above the ground. This was done for 21 seconds per quarter (1/4 of 84 s being the total application time per plot) and then repeated for the other quarters of the same field plot. To avoid contamination of surrounding plots, each plot was enclosed by a 2 m high plastic sheet during application.

#### 4.2.3. Exposure analysis

Seven days after insecticide application, vegetation samples were collected from inside the field plots to quantify the amounts of the active ingredient acetamiprid and the metabolite acetamiprid-N-desmethyl by LC-MS analysis. This was done to demonstrate the potential presence of the insecticide inside the enclosures with respect to the different treatments even days after application as well as its metabolization within plant the tissue. The full analytical procedure and results are given in supplementary file 1.

#### 4.2.4. Insect sampling and identification

Insects in the upper vegetation of each field plot were regularly sampled with a sweep net (Flexi-Kescher Dreieck; Bioform, Nürnberg, Germany). Sampling took place 2, 9, 16, 24 and 30 days after insecticide application. While walking parallel to the plot, the net was swept four times forwards and backwards across the upper part of the vegetation. After the fourth sweep, the net was immediately closed by hand. The entire contents were then transferred into a three-litre zip-lock plastic bag (Profissimo, dm-drogerie markt GmbH + Co. KG, Karlsruhe, Germany). The plastic bags were stored at -20 °C for at least 24 hours to kill all arthropods inside.

For evaluation, individual bags were stored at 8 °C for 30 min to carefully unfreeze the specimens. The entire contents of the plastic bag were then transferred to a plastic tray and the plant bugs (imagines and nymphs) were separated with tweezers. Nymphs were counted and transferred to a 2 ml screw-capped microtube (SARSTED AG & Co. KG, Numbrecht, Germany) filled with 70% ethanol for preservation. Imagines were sorted by morphospecies, and the corresponding individuals were counted. For each morphospecies, one male and one female specimen (if available) were separated for preparation. The specimens were mounted on paper cards using wallpaper glue. In the case of male specimens, the aedeagi were removed beforehand and glued next to the respective specimen. Remaining specimens of each morphospecies were preserved in 70% ethanol as described above. All morphospecies were identified to species level using the Hackston (2023) Miridae key and the Corisa software (Strauss and Simon 2021).

#### 4.2.5. Statistical evaluation

A generalised linear mixed model (GLMM) was fitted for the combined individual counts of nymphal and adult plant bugs using the ‘lme4’ package (version 1.1-33, Bates et al. 2015) with ’*count*’ as the explanatory variable, ’*treatment*’ as a fixed effect, ’*sampling*’ as a co-factor, and ’*block*’ and ’*plot*’ as random effects assuming a Poisson distribution. An interaction between treatment and sampling was included. Model fit was assessed by residual analysis, considering the R2 value and the Akaike Information Criterion (AIC), and by checking for possible over-dispersion. Significant factors and interactions within the model were identified using a post-model ANOVA. Those were ’*treatment*’, ’*sampling*’ and the interaction between both factors. Pairwise post hoc multiple comparisons by sampling were then executed using the Tukey-test based on estimated marginal means using the emmeans package (Lenth 2023) considering the *treatment*-*sampling* interaction.

In a second step, a separate GLMM was fitted including only the count data of *S*. *binotatus*, *L*. *dolabrata* and *M*. *recticornis* and only the data of the first sampling (i.e., two days after insecticide application). In this GLMM, ’*count*’ was the explanatory variable with ’*treatment*’ as a fixed effect, ’*species*’ as a co-factor, an interaction between ’*treatment*’ and ’*species*’ and ’*block*’ and ’*plot*’ as random effects, assuming a Poisson distribution. Model fit was assessed as described above. Significant factors and interactions within the model were identified using a post-model ANOVA. Pairwise post hoc multiple comparisons by species were then executed using the Tukey-test based on estimated marginal means considering the treatment-species interaction.

Based on the control sampling data of the three species, the populations were stable in the early samples, but declined dramatically in later samples as expected based on the known phenologies. In order to compare acute species-specific treatment response, we therefore did not include data beyond the first sampling. The complete R-code is provided in supplementary file 5.

### 4.3. Greenhouse Experiment

#### 4.3.1. Host plant preparation

Eighteen plots of 30 cm diameter with live plants, consisting mainly of grasses, which are the host plants of the three mirid species, were collected from a meadow with a shovel and transferred to 12-liter plant pots. These were previously filled halfway with a 50:50 mixture of sand and humus. Only sections of the meadow with a high proportion of grasses were selected to provide sufficient host plants for the plant bugs. The pots were placed in an outdoor test facility for one month and watered as needed until the grasses had developed seed heads.

#### 4.3.2. Insecticide application and experimental procedure

The pots were placed in an 8 m^2^ rectangular area, surrounded by 2 m high vertical plastic sheathing. Spray application was carried out using the same protocol and equipment as described for the field experiment. Two treatments were applied: a solution containing 30% field rate concentration of Mospilan®SG and a control containing only water. For each treatment application, nine randomly selected plant pots were placed within the application area, all 30 cm apart. The control was applied first. The pots were then placed in the outdoor test facility. A total of four blocks were formed, with two blocks for each treatment, one containing five pots and the other one containing four. Treatment and control blocks with the same number of replicates were placed opposite to each other (Fig. 5). In the center of each pot, a 2 m wooden stick was vertically inserted. At the top of the stick, a 30 cm diameter herbacaeus ring was fixed horizontally. Fine insect gauze was then placed over the construction from the top of the stick down to the plant pot, forming a closed tube around the host plants.

**Figure 5:**
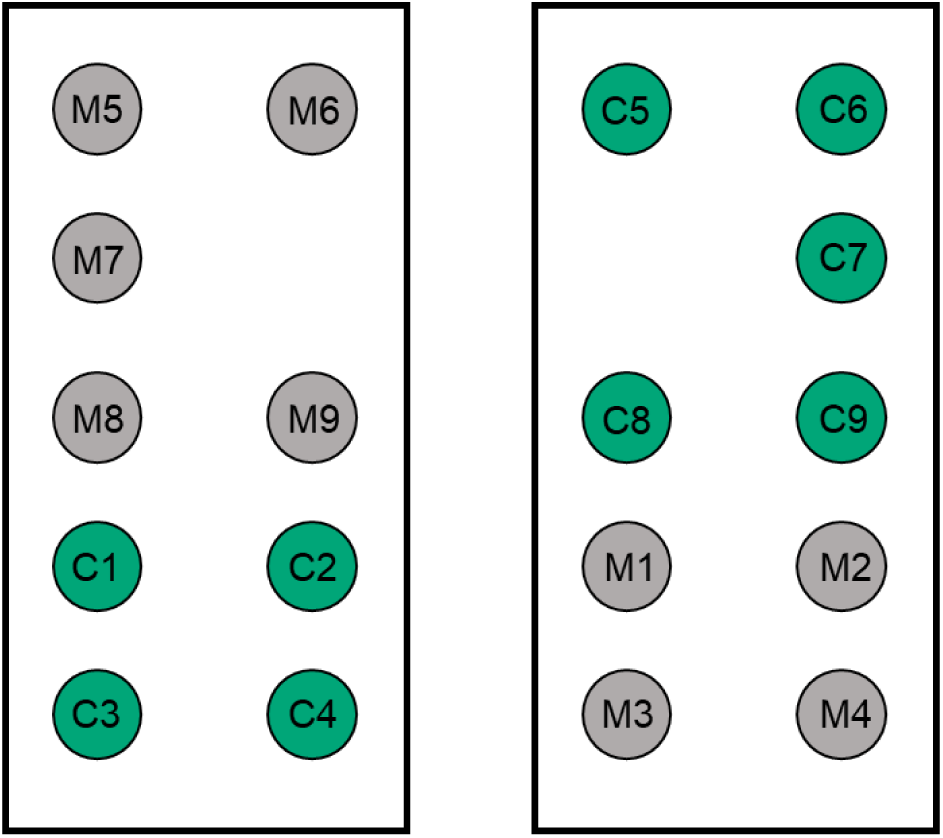
Schematic representation of the experimental setup. "M"=Mospilan®SG treatment, "C"=Control treatment. Treatment and control blocks with the same number of replicates are placed opposite to each other.

Two days after application, plant bugs were caught using a butterfly sweep net (Flexi-Kescher Dreieck; Bioform, Nürnberg, Germany) in the same extensively managed meadow from which the plant samples were collected. Specimens were immediately transferred to 30 cm x 30 cm AERARIUM mesh cages (Bioform, Nürnberg, Germany), which were then placed in an 8 °C room. After 15 min of settling, the cages were opened and specimens of *S*. *binotatus*, *L*. *dolabrata* and *M*. *recticornis* were sorted by species (identified by naked eye) and transferred to petri dishes. For each replicate (= one plant pot), three petri dishes were prepared, each containing six specimens of either species (males and females mixed). After assuring that all bugs were vital, 18 plant bugs (6 per species) were then released on top of the plants inside the insect gauze. At the same time, plant samples for HPLC analysis were collected from each Mospilan®SG treated plot to demonstrate that the active ingredient acetamiprid was present at the time of plant bug release. Each morning the plants were sprayed with a pressurized water bottle to simulate morning dew. After four days (six days after insecticide application) the experiment was terminated, and alive specimens were counted. Specimens from each replicate were stored separately in 2 ml micro tubes with screw caps (SARSTEDT AG & Co. KG, Germany) filled with 70% ethanol. Following the assay, species identity was again verified using a binocular and the Hackston (2023) Miridae key and the Corisa software (Strauss and Simon 2021).

#### 4.3.3. Exposure analysis

Two days after insecticide application, just before the plant bugs were released into the cages, vegetation samples were collected from each replicate of the insecticide treatment group. Unlike the field experiment, the control group was not sampled as the control pots were sprayed first and then removed before insecticide application. LC-MS analysis was used to quantify the amounts of the active ingredient acetamiprid and the metabolite acetamiprid-N-desmethyl. The full analytical procedure and results are provided in supplementary file 1.

#### 4.3.4. Statistical evaluation

A linear mixed regression model, was fitted using the ‘lme4’ package (Bates et al. 2015) with ’*alive*’-specimens as the response variable, ’*treatment*’ as fixed effect and ’*species*’ as a co-factor, and an interaction between ’*treatment*’ and ’*species*’. The random factor ’*block*’ was first included but did not explain any variance of the data and was therefore excluded from the model. The random factor ’*pot*’, providing a unique identifier for each pot, was included to account for the fact that ’*species*’ was nested within ’*treatment*’, allowing an individual baseline morality for each replicate. Hence, if the interaction was significant, it was not due to a replicate effect, allowing to explore species differences in survival only. Significance of the interaction between ’*treatment*’ and ’*species*’ was confirmed by a post model ANOVA.

Fitting a follow-up model with ’species’ as the explanatory variable was not possible because there were no individuals of *S*. *binotatus* that survived the insecticide treatment in any of the nine replicates, making it impossible to estimate a standard error. Accordingly, we report the difference in treatment effect as the percentage difference in survival rates between the control and insecticide treatments (’insecticide effect’). This considers the baseline mortality of each species observed in the control.

The complete R-code is provided in supplementary file 5.

### 4.4. Dose-Response Assays

#### 4.4.1. Insecticide preparation

A stock solution of 4.16 mg/ml Mospilan®SG, dissolved in tap water, was prepared corresponding to 10 times the highest recommended field rate of 0.416mg/ml. Stock and treatment solutions were stored in 1.5 ml ND8 amber glass screw-neck vials (LABSOLUTE – TH GEYER) in the dark at 4°C. Stock solutions were renewed weekly. Prior to each dose-response assay, a dilution series was prepared based on the stock solution on the day of application (Tab. 3).

**Table 3:** Treatments applied in the dose-response assay and respective Mospilan®SG concentrations. The 0.14% treatment was not applied to *Leptopterna dolabrata* and *Megaloceroea recticornis* due to lower sensitivity.

| Treatment | Concentration [ng/μl] |
| --- | --- |
| Control | 0.00 |
| 100% | 416.00 |
| 33.30% | 138.67 |
| 11.10% | 46.22 |
| 3.70% | 15.41 |
| 1.24% | 5.14 |
| 0.41% | 1.71 |
| 0.14% | 0.57 |

Here, 0.5% TWEEN (20 was added to the solvent to break the surface tension and ensure easy application to the bug’s body surface. Each treatment concentration was prepared from the stock solution by pipetting volumes from higher to lower concentration treatment tubes prefilled with pure solvent (see dilution protocol supplementary file 6).

#### 4.4.2. Insect sampling

Plant bugs were caught in the meadows of the Hohenheim Gardens (extensively managed grassland) using a butterfly sweep net (Bioform, Nürnberg, Germany). Specimens were immediately transferred to 30x30 cm insect mesh cages (Bioform, Nürnberg, Germany), which were then placed in an 8 °C room. After 15 min of settling, the cages were opened and specimens of *S*. *binotatus*, *L*. *dolabrata* and *M*. *recticornis* were sorted by species (identified by naked eye) and transferred to petri dishes. Two petri dishes were prepared per species and treatment with six to four specimens each (depending on availability of specimens). Where possible, male and female specimens were equally distributed. Only vital specimens that were actively climbing the net were included. Petri dishes were taken to the laboratory (20°C) where the specimens were allowed to settle again for 20 min. Prior to treatment application, all specimens were again checked for vitality and replaced if necessary.

#### 4.4.3. Topical application and evaluation

Several identical assays were run in sequence to allow for a possible cohort effect (*S*. *binotatus* + *M*. *recticornis*: three assays; *L*. *dolabrata*: four assays). Each assay included two physical replicates per concentration, each consisting of four to six specimens (2 male + 2 female / 3 male + 3 female) based on availability of specimens in the Hohenheim Gardens. The specimens of a physical replicate were first placed in a petri dish and later transferred to rearing boxes (see below). Accordingly, each concentration consisted of six-(*S*. *b*. + *M*. *r*.) or eight replicates (*L*. *d*.) nested within three or four cohorts (Tab. 4). Note that for *M*. recticornis the number of replicates in the lowest tested Mospilan®SG concentration (0.41%) is four. The number of treatments for *S*. *binotatus* was increased for the second and third cohort adding the 0.14% treatment due to higher sensitivity.

**Table 4:** Number of cohorts, treatments and individuals tested in the dose-response assays. Eight treatments: 100% - 0.14% + control; seven treatments: 100% - 0.41% + control. Note that the number of treatments for *S*. *binotatus* was increased for the second and third cohort.

| Species | Cohort | Number of treatments | Individuals tested per treatment |
| --- | --- | --- | --- |
| *S. binotatus* | 1 | 8 | 4 |
|  | 2 | 8 | 6 |
|  | 3 | 8 | 6 |
| *L. dolabrata* | 1 | 7 | 6 |
|  | 2 | 7 | 6 |
|  | 3 | 7 | 6 |
|  | 4 | 7 | 6 |
| *M. recticornis* | 1 | 7 | 4 |
|  | 2 | 7 | 6 |
|  | 3 | 7 | 6 |

Bugs inside petri dishes were anaesthetized with CO2, then 1 µl of insecticide solution (and pure solvent as a control) was applied dorsally to each specimen. Application was conducted using the Hamilton repeating dispenser with a 50 µl Hamilton syringe (Hamilton Bonaduz AG, Switzerland). Specimens of each replicate were then transferred to 500 ml plastic cups. The cups contained a water supply (Eppendorf tubes filled with water and cotton wool), a bundle of grasses from the original habitat with the top 10-15 cm of the plants cut off (stems and heads) placed in an Eppendorf tube filled with water and cotton wool, and a paper tissue covering the bottom of the cup. The lid was provided with several air holes. In addition, water was sprayed into the cup to cover the grasses with droplets providing sufficient humidity and an additional source of drinking water. Cups were stored under standard laboratory conditions. After 24 h, mortality rates for all treatments were recorded for each replicate and sex. The experiment was then terminated and dead and surviving specimens from each replicate were stored separately in 2 ml micro tubes with screw caps (SARSTEDT AG & Co. KG, Germany) filled with 70% ethanol. Following the assay, species identity and sex of all specimens were again verified using a binocular and the Hackston (2023) Miridae key and the Corisa software (Strauss and Simon 2021).

#### 4.4.4. Dose-response analysis

Data from each assay were first analyzed with a generalized linear mixed model using the ‘lme4’ package (Bates et al. 2015). ’*Mortality*’ was included as the explanatory variable, ’c*oncentration*’ as fixed effect, ’*sex*’ as co-factor and ’*replicate*’ and ’*cohort*’ as random effects. Additionally, an interaction between ’c*oncentration*’ and ’*sex*’ was included. Model fit was assessed by residual analysis, considering the R2- and AIC value, and by checking for possible over-dispersion. Including the random effect ’*replicate*’ allowed to control for the fact that ’*sex*’ was nested within ’*concentration*’, allowing an individual baseline morality for each replicate. If the interaction was significant, it was not due to replicate effect, allowing to generate LD_50_ values for male and female specimens of the same species separately.

A post-model ANOVA was performed confirming that ’*concentration*’ was a significant predictor in all assays. A significant interaction between ’*concentration*’ and ’*sex*’ was found for *L. dolabrata* and *M. recticonis*.

Log-logistic dose-response models were then fitted for a combined species analysis and for each species while modelling male and female response separately using the ’drc’ package (Ritz et al. 2015). In the case of control mortality, the lower asymptote of the fitted regression curve was not set to ’0’ to generate unbiased parameter estimates. The upper asymptote was set to ’1’. The best model fit was selected based on visual inspection of the regression line and evaluation of the confidence intervals of the model parameters, considering the AIC estimator. Dose-response curves were generated using the plot() operation. LD_50_ values were obtained using the ED() function, including a control for possible over-dispersion of the logistic regression model. Differences between LD_50_ values of individual curves within the same model were analyzed using the compParm() function executing approximate t-tests with α = 0.05 and including a correction for possible over-dispersion of the underlying model.

The complete R-code for the linear and log-logistic models is provided in supplementary file 5.

## Supporting information

Supplementary file 1_pesticide exposure analysis

Supplementary file 2_fresh weight & weight normalized LD50

Supplementary File 3_Environmental Conditions Field Experiment

Supplementary File 4_Calibration of the Field Compression Sprayer

Supplementary file 5_statistical report

Supplementary file 6_Mospilan(R)SG dilution protocol

## Acknowledgements

We would like to thank Sarah Rißmann for her constant support throughout this project and Nina Estler for her help with the dose-response assays. Further, we would like to thank Denise Kuhn, Anja Betz, Robert Bischoff and Prayan Pokharel for their support and critical feedback. Special thanks go to Manfred Konert for his help in setting up and maintaining the field experiment. In addition, we would like to thank the team at the Botanical Garden Hohenheim, led by Dr Helmut Dalitz, for their support in carrying out the field trial. We are also grateful to Jürgen Deckert, who advised us on the subject of soft bug handling, and to Gerhard Strauss for the permission to use insect images from the Corisa software. Finally, we would like to thank the Graduate Academy of the University of Hohenheim, which made this project possible by providing a scholarship as part of the Baden-Württemberg Graduate Support Programme.

## Contributions

CA.B., I.G. and G.P. supervised the project. JE.S., CA.B., I.G. and G.P. drafted the experiments. JE.S. and P.B. conducted the field experiment. JE.S. performed the greenhouse experiment. JE.S. performed the dose-response assays. JE.S. performed the analysis of the field- and greenhouse experiments. JE.S., D.G. and HP.P. performed the analysis of the dose- response assays. B.H., F.W. and JE.S. conducted the pesticide analysis. JE.S. wrote the original draft of the manuscript. G.P., CA.B., I.G., HP.P. edited the manuscript.

