## Supplementary file 1_pesticide exposure analysis for "The neonicotinoid acetamiprid is highly toxic to wild non-target insects"

**Protocol of the Pesticide Exposure Analysis of Plant Material**

**–**

**Field- and Greenhouse Experiment**

### Sampling - field experiment

Composite vegetation samples were collected from the interior of all plots at 7-, 17-, 30-, and 60 days after insecticide application. At 7- and 17 days after application, six individual samples were taken inside every enclosure by cutting all above-ground plant material at a 10 x 10 cm area. At 30- and 60 days after application, the number of individual samples was reduced to three to minimize the impact on the vegetation inside the enclosures. Plant material from the same field enclosure was mixed and placed in a plastic bag and stored at -20°C.

### Sampling – greenhouse experiment

Two days after insecticide application (30% field rate Mospilan®SG), host plant material (grasses) was sampled from the insecticide-treated cages only. In each cage, two blades of grass were sampled. The grass was cut directly above the ground with a pair of scissors. Plant material from the same cage was directly transferred to a plastic bag and stored at -20 °C.

### Extraction

Frozen plant material from each sample was placed in a glass dish, cut into 2 cm pieces and thoroughly mixed. Then, 2 g (wet weight) of this plant material was crushed in a mortar with the addition of liquid nitrogen to allow the material to macerate easily. The sample was then transferred to a 100 ml centrifuge tube. Acetonitrile (20 ml) (ACN) and 250 µl of distilled water were added. The sample was homogenised for 1 min using a Miccra MiniBatch D-9 (MICCRA GmbH, Heitersheim, Germany). The contents were filtered into a 25 ml volumetric flask using filter paper. ACN was added to give a total volume of 25 ml. After mixing, 2 ml were transferred to a 2 ml Eppendorf tube and 20 µl ethylene glycol was added. The sample extracts were reduced to 200 µl in the vapotherm mobil s (Barkey GmbH & Co KG, Leopoldshöhe, Germany) at 60 °C under a constant flow of nitrogen gas. The remaining volume was filled up to 0.5 ml with an ACN/water mixture (3:7) and then centrifuged at 12000 rpm for 10 min (miniSpin, Eppendorf AG, Hamburg, Germany). The supernatant was transferred to auto sampler glass vials using a syringe with a 0.45 µm PTFE filter (ROTILABO® Mini-Tip, Carl Roth GmbH + Co. KG, Karlsruhe, Germany).

**Quantification**

To quantify acetamiprid and acetamiprid-N-desmethyl (metabolite of ACE), 3 µl of the extract was injected into an LTQ Velos LC-MS system (Thermo Fischer Scientific, Waltham, Massachusetts, USA) and the compounds were separated on a Kinetex® 2.6 µm XB-C18 100 Å, 150 x 4.6 mm, LC column (Phenomenex Ltd. Deutschland, Aschaffenburg, Germany). Compounds were eluted at 40 °C at a constant flow rate of 0.5 ml/min using the following water + 10% acetonitrile (A) - acetonitrile (B) gradient: 0-1 min 95% A, 10 min 40% A, 12-14 min 10% A, and 14.1-16 min 90% A. Compounds were detected using acetamiprid/acetamiprid-N-desmethyl standards at 0.005, 0.01 and 0.1 mg/l.

To calculate the amount of compound per gram of plant tissue, the dry weight of each plant sample was determined to correct for the wet weight (see Extraction). For this purpose, the same amount of plant tissue (wet weight) used for extraction (2 g) was placed in a previously weighed glass petri dish and stored at 80 °C in a Heraeus T12 oven (Heraeus Instruments, Hanau, Germany) for 24 h. The petri dish was weighed again to determine the dry weight.

The following formula was used to determine the amount of compound per gram of plant tissue (dry weight):

$$amount\ in\ \frac{\mu g}{g} = \frac{X\ \mu g/ml * 0.5\ ml * 25\ ml/2\ ml}{sample\ weight\ (dry)\ in\ g}$$

### Results – Field experiment

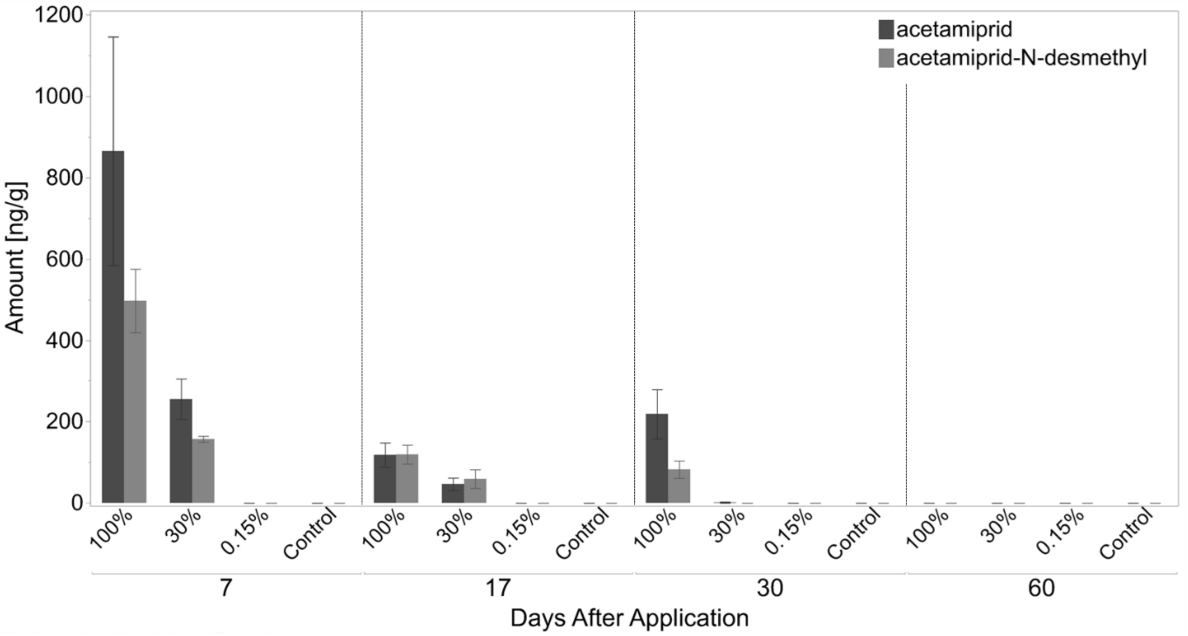


**Figure 1: Mean amount of acetamiprid and the metabolite acetamiprid-N-desmethyl (± SE) per gram plant tissue sampled inside field enclosures (LC-MS)** seven-, seventeen-, thirty- and sixty days after application of Mospilan®SG treatments and the control (water). ND: not detected; n = 5 for each treatment and

**Table 1: Mean amount of acetamiprid and the metabolite acetamiprid-N-desmethyl (± SD) per gram plant tissue sampled inside field enclosures (LC-MS)** seven-, seventeen-, thirty- and sixty days after application of Mospilan®SG treatments and the control (water). Round numbers are given, n = 5 for each treatment and substance.

| | **Treatment** | **Mean Amount Day 7 [ng/g]** | **Mean Amount Day 17 [ng/g]** | **Mean Amount Day 30 [ng/g]** | **Mean Amount Day 60 [ng/g]** |
|---|---|---|---|---|---|
| **acetamiprid** | 100% | 865 (± 628) | 118 (±66) | 219 (±135) | 0 |
| **acetamipird-N-desmethyl** | 100% | 497 (±175) | 119 (±52) | 81 (±47) | 0 |
| **acetamiprid** | 30% | 255 (±111) | 46 (±34) | 1 (±3) | 0 |
| **acetamipird-N-desmethyl** | 30% | 157 (±16) | 59 (±51) | 0 | 0 |

### Results – Greenhouse Experiment

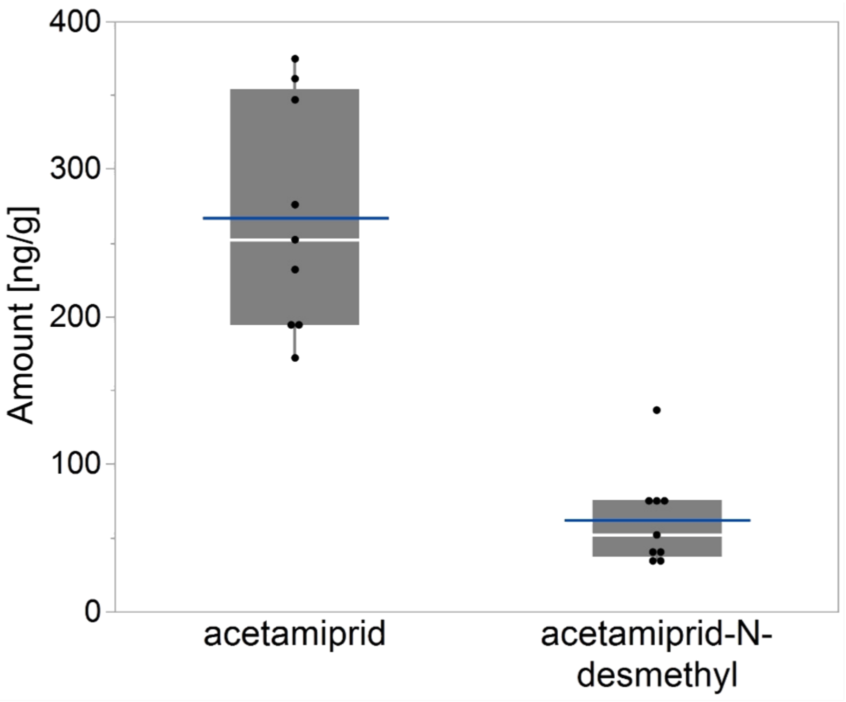


**Figure 2: Amount of acetamiprid and the metabolite acetamiprid-N-desmethyl in plant tissue** sampled inside cages two days after spray application of 30% Mospilan®SG (LC-MS). Blue horizontal line indicates mean value (acetamiprid: 267 ng/g, acetamiprid-N-desmethyl: 62 ng/g); n = 9 for each substance.
