## Supplementary File 3_Environmental Conditions Field Experiment for "The neonicotinoid acetamiprid is highly toxic to wild non-target insects"

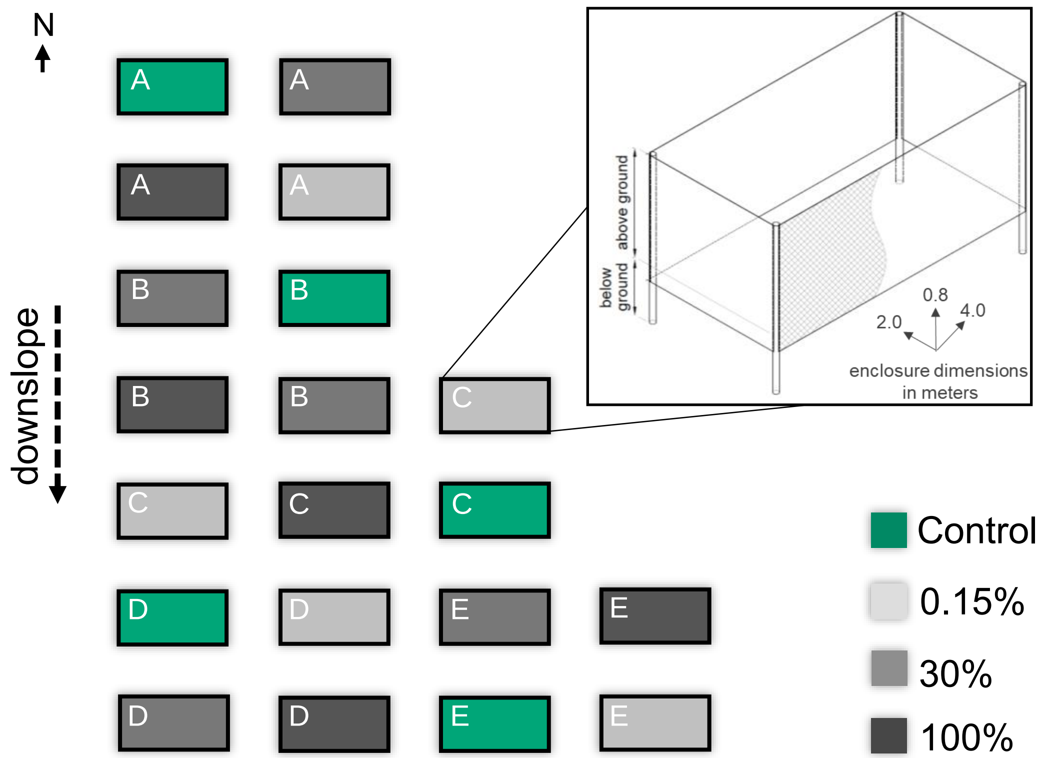


**Figure 1: Environmental conditions in the field trial.** Field enclosures are separated by 2 m. Coloration indicates Mospilan®SG – treatment based on contamination estimates. Drawings created with AUTODESK AutoCAD 2023 and Fusion 360. White letters inside plots indicate the assigned block. Drawings created with AUTODESK AutoCAD 2023 and Fusion 360.
