## Supplementary File 4_Calibration of the Field Compression Sprayer for "The neonicotinoid acetamiprid is highly toxic to wild non-target insects"

**Calibration procedure of the compression sprayer used in the Field- and Greenhouse Experiment**

To ensure that the exact amount of 0.48 l per field plot is applied, the hand sprayer was calibrated beforehand. Here, we considered several factors: without a manometer, the pressure must be set to an equal level before every spraying event. Therefore, the number of pumps to increase the pressure inside the bottle was set to 40. Additionally, a certain amount of residue liquid remains in the pressure bottle after every spraying event. Therefore, the following calibration procedure was performed.

- Water (0.48 l) was filled into the empty pressure bottle. After 40 pumps, the spray gun was opened until no more liquid exited the nozzle.
- The pressure bottle was opened and 0.48 l of water was added on top of the residue after the first spraying. The pressure was again increased by 40 pumps and the spray gun was opened over a measuring cylinder while recording the time. As soon as the first air from inside the pressure bottle exited the nozzle, the spray gun was closed. The time was stopped and the amount of liquid inside the measuring cylinder was noted.
- This procedure was repeated several times. The mean time for spraying was 84 s and the mean deviation of liquid input and output was about 1% (input: 0.48 l, output: 0.475 l).
- Accordingly, the time of spraying for every field plot was set to 84 s.

To control for an unequal application due to a change in volumetric flow rate (as a result of decreasing pressure during spraying), the field plots were subdivided into four equal quarters. Every quarter was given a number from one to four. The spraying would always start at the quarter with number one and end at number four. The arrangement of numbers was then randomized within plots of the same treatment. Based on the total application time per plot, the application time per quarter was 21 seconds.
