## Supplementary file 6_Mospilan(R)SG dilution protocol for "The neonicotinoid acetamiprid is highly toxic to wild non-target insects"

**Mospilan®SG Dilution Protocol – Dose-Response Assays**

Field rate of Mospilan®SG: 250 g/ha in 200-600 l/ha -> Target: 1 µl/insect topical application

$$\frac{250\ g}{600\ l} = 0{,}416\ \frac{g}{l} = 0{,}416\ \frac{mg}{ml}$$

Prepare stock solution (10x 100%-Sol.):

$$100\% - Sol, equals \rightarrow \frac{0{,}2\ g}{0{,}48\ l} = \ 0{,}416\ \frac{g}{l} = 0.416\ \frac{mg}{ml}$$

$$Stock\ soultion\ should\ be \rightarrow 4{,}16\ \frac{mg}{ml}\ (10x\ 100\% - Sol,)$$

Weighing example:

$$\frac{weighed\ portion\ mg}{dilute\ in\ Xml} = \frac{4{,}16\ mg}{1\ ml} \rightarrow Xml = \frac{weighed\ portion\ mg}{4{,}16\ \frac{mg}{ml}}$$

$$100\ \mu l\ (10x\ stock) + 900\ \mu l\ water\ =\ 1\ ml\ 100\% - Sol,$$

$$\downarrow\ - 250\ \mu l\ (750\ \mu l\ of\ 100\%\ left)$$

$$250\ \mu l\ (100\% - Sol,) + 500\ \mu l\ water = 750\ \mu l\ 33{,}3\% - Sol,$$

$$\downarrow\ - 250\ \mu l\ (500\ \mu l\ of\ 33{,}3\%\ left)$$

$$250\ \mu l\ (33{,}3\% - Sol,) + 500\ \mu l\ water = 750\ \mu l\ 11{,}1\% - Sol,$$

$$\downarrow\ - 250\ \mu l\ (500\ \mu l\ of\ 11{,}1\%\ left)$$

$$250\ \mu l\ (11{,}1\% - Lsg) + 500\ \mu l\ water = 750\ \mu l\ 3{,}7\% - Sol,$$

$$\downarrow\ - 250\ \mu l\ (500\ \mu l\ of\ 3{,}7\%\ left)$$

$$250\ \mu l\ (3{,}7\% - Sol,) + 500\ \mu l\ water = 750\ \mu l\ 1{,}24\% - Sol,$$

$$\downarrow\ - 250\ \mu l\ (500\ \mu l\ of\ 1{,}24\%\ left)$$

$$250\ \mu l\ (1{,}24\% - Sol,) + 500\ \mu l\ water = 750\ \mu l\ 0{,}41\% - Sol,$$

$$\downarrow\ - 250\ \mu l\ (500\ \mu l\ of\ 0{,}41\%\ left)$$

$$250\ \mu l\ (0{,}41\% - Sol,) + 500\ \mu l\ water = 750\ \mu l\ 0{,}14\% - Sol,$$

Vortex in between all steps!
